# Deficiency of *Shank3* in the Nucleus Accumbens Reveals a Loss of Social-Specific Motivation

**DOI:** 10.1101/2025.01.08.631742

**Authors:** Oakleigh M. Folkes, Meaghan Donahue, Sung Eun Wang, Nicole Xinyen Oo, Sheng-nan Qiao, Xiaoming Wang, Paola N. Negrón-Moreno, Yong-hui Jiang

## Abstract

Deficits in social interaction are a hallmark symptom of autism and other neuropsychiatric disorders. *SHANK3* encodes a postsynaptic density scaffold protein and is one of the most common causal genes for autism. SHANK3 protein is highly expressed in the nucleus accumbens (NAc), a critical brain region underlying motivated behavior, including social motivation. We previously reported that global *Shank3^Δe^*^4–22^ deletion mice have decreased motivation for palatable food, increased unilateral social investigation, and show a hypoactive NAc and NAc-connected circuits. We thus developed a new *Shank3^flox/flox^*mouse tool to conditionally knockdown SHANK3 in a region-specific manner. We found that knockdown of *Shank3* in the NAc decreased social preference in the 3-chamber assay and decreased social motivation in the social conditioned place preference (sCPP) assay. *Shank3-*NAc deletion did not alter food reward seeking, reciprocal social investigation, or anxiety-like behaviors, that we report in global *Shank3^Δe^*^4–22^ deletion mice. These data establish a novel and specific role of *Shank3* in the NAc on social motivation.

## INTRODUCTION

Deficits in social interaction are a hallmark symptom of autism spectrum disorder (ASD) and other neuropsychiatric disorders.^1^ ASD genomic studies have identified a large number of genes that encode proteins involved in synaptic development and functions, suggesting a potential converging mechanism.^2–13^ *SHANK3* encodes a postsynaptic density scaffold protein and is one of the most common causal genes for autism, found in ∼2% of patients, and has been extensively modeled in animals.^14–16^

To study the role of SHANK3 on synaptic function and behavior, we previously developed the *Shank3* complete deletion model with an exon 4 to 22 deletion (*Shank3^Δe^*^4–22^).^17^ We showed that *Shank3^Δe^*^4–22^ mice have increased unidirectional social behavior in a naturalistic setting, are profoundly impaired in reward-seeking behavior in a lever press task for food pellets, and display anxiety-like behavior compared to wild-type (WT) controls.^17^ These data corroborate extensive literature showing that removing *Shank3* isoforms impacts social behaviors, indicating SHANK3 expression is a fundamental piece of neural mechanisms of social interaction.^18–24^ Yet, how *Shank3* in specific brain regions coordinates social behaviors, motivation, and anxiety-like behavior is not well understood.

SHANK3 protein is highly expressed in the nucleus accumbens (NAc),^24–27^ a well-established regulator of reward, social interaction, and social motivation.^28–34^ Imaging studies in autistic individuals reveal altered NAc responses to social cues,^35–37^ and many autism mouse models show critical NAc activity changes,^38–44^ indicating it is an essential region for regulating social behaviors relevant to autism. We previously demonstrated that the NAc and NAc neuronal networks have a blunted response to social stimuli in *Shank3^Δe^*^4–22^ mice,^17^ and others have shown that removing *Shank3* only from the NAc is sufficient to drive lower NAc firing rates.^27^ This supports the hypothesis that SHANK3 in the NAc is critical for driving the behavioral and neural deficits in *Shank3^Δe^*^4–22^ mice and highlights the importance of the NAc as a central hub for the loss of social and reward-seeking behaviors in *Shank3^Δe^*^4–22^ mice. Yet, investigation into the role of SHANK3 in the NAc on social behavior is limited.

Here, we show that removing *Shank3* from the NAc display normal anxiety-like and locomotion behaviors in open field and selectively impairs social motivation but does not alter food motivation. This starkly contrasts with conventional *Shank3^Δe^*^4–22^ mice, which strongly prefer a socially-paired chamber after conditioning, have a complete deficit of food reward-seeking, and anxiety-like and hypo-locomotion behaviors. Our data suggest that *Shank3* in the NAc underlies social motivation independent from other reward-seeking behaviors.

## RESULTS

### Conditional Deletion of *Shank3*^Δe^^4–22^ from the Nucleus Accumbens Significantly Reduces SHANK3 Levels in the Synaptosome

Previously, we established a conditionally floxed *Shank3*^Δe4–22^ *^fl/fl^* line with three loxP sites in intron 3, 9, and 3′ UTR after exon 22^45^ that showed a mixture of recombination between exon 4-9 and exon 4-22 leading to variability in *Shank3* isoform expression in this model. We generated a new *Shank3*^Δe4–22^ mouse model with two loxP sites flanking exon 4-22 that eliminated the possibility of competition among 3 loxP sites (**Fig. 1A, S1 A-C)**.

**Figure 1:**
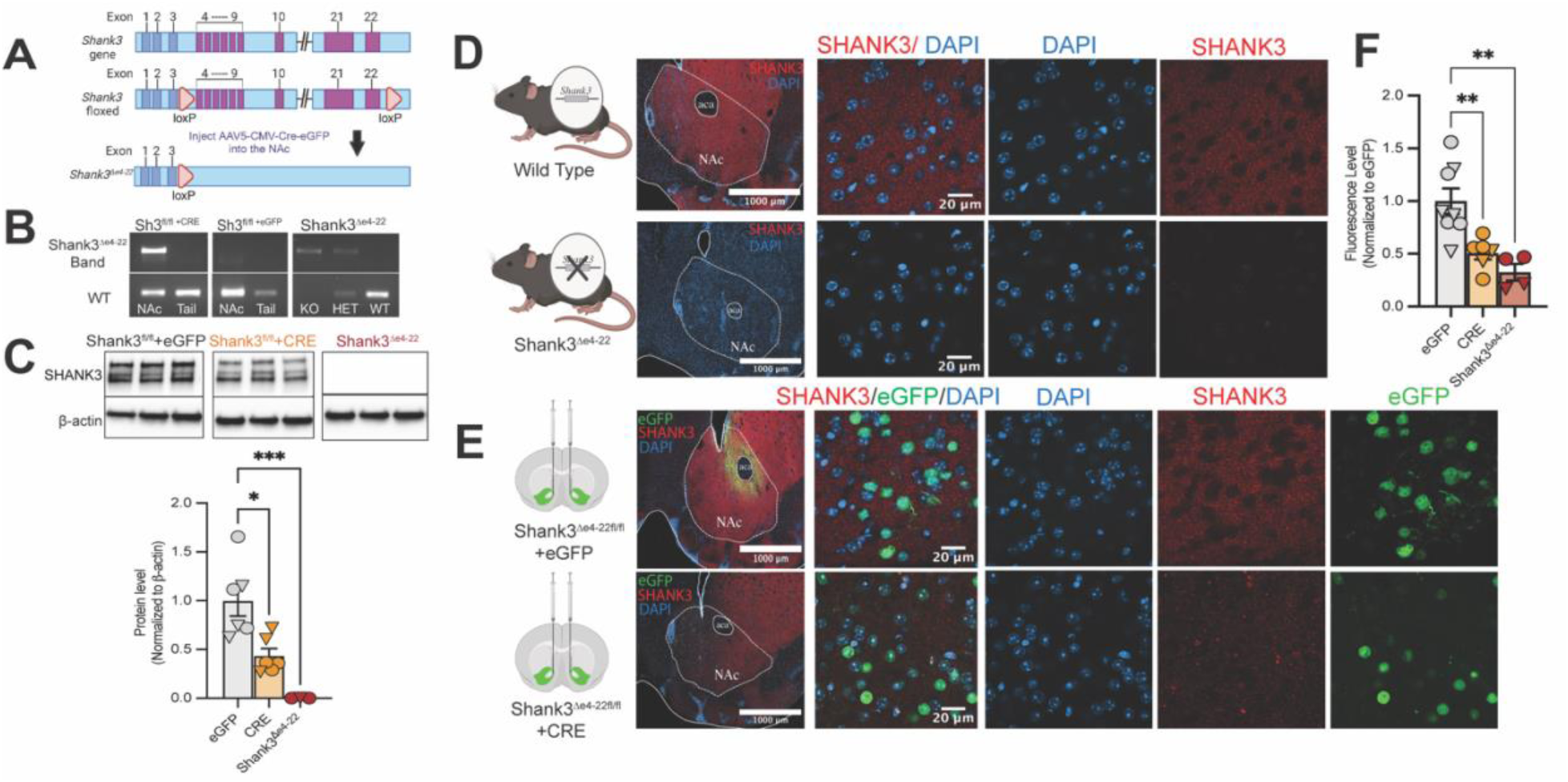
Brain Region-Specific Knockdown of *Shank3* Reduces SHANK3 protein expression in the NAc. **A)** Schematic design of *Shank3* deletion strategy using CRE*-loxP*. Arrows indicate *loxP* sites. Exons removed are shown in purple. AAV5-CMV-Cre-eGFP was injected directly into the NAc of *Shank3^fl/fl^* animals to delete Δe4-22 of *Shank3*. **B)** PCR revealed the detection of *Shank3* deletion e4-22 in the NAc but not the tail in *Shank3^fl/fl^* mice injected with *Cre* and *Shank3^Δe^*^4–22^ controls. **C)** SHANK3 protein levels were quantified using western blot analysis in eGFP or Cre-injected *Shank3^fl/fl^*mice and *Shank3^Δe^*^4–22^ mice. SHANK3 is significantly decreased in Cre injected *Shank3^fl/fl^* mice (*\*P=0.0123*) and *Shank3^Δe^*^4–22^ mice (*\*\*\*P=0.0008)* compared to eGFP controls (Ordinary one-way ANOVA, *F_(2,12)_=14.25; P=0.0007; n1=6, n2=6, n3=3*). **D-F**) Immunostaining of SHANK3 (red) and DAPI (blue) with eGFP (green) from viral injection. Representative 20x and 63x images of the NAc with SHANK3 immunostaining showed **D**) SHANK3 was absent in *Shank3^Δe^*^4–22^ mice and **E**) reduced in *Shank3^fl/fl+CRE^* mice compared to *Shank3^fl/fl+eGFP^* mice. **F**) A secondary antibody fluorescent signal was used to visualize the SHANK3 primary antibody. Fluorescence was decreased significantly in *Shank3^fl/fl+CRE^* mice compared to *Shank3^fl/fl+eGFP^*(** *P= 0.0078*) and in *Shank3^Δe^*^4–22^ mice compared to *Shank3^fl/fl+eGFP^* (\*\**P=0.0018*) (Ordinary one-way ANOVA, *F_(2,15)_=11.28; P=0.0010; n1=8, n2=8, n3=6*). Complete statistical analyses are provided in the statistical analyses file associated with this manuscript.

We conditionally deleted *Shank3* exclusively from the NAc using a viral-mediated approach. AAV5-CMV-Cre-GFP or AAV5-CMV-eGFP control was injected into the NAc of adult male and female *Shank3^fl/fl^* mice. To assess the recombination efficacy between loxP sites, we first took NAc brain punches and conducted PCR to identify the presence of joint fragments between loxP sites **(Fig. 1B)**. We did not see recombination in *Shank3^fl/fl^* mice injected with eGFP. We next confirmed our approach of genetically removing *Shank3* was sufficient to reduce protein expression. We found that the expression of SHANK3 protein in *Shank3^fl/fl^* mice injected with CRE was reduced by approximately 50 percent of controls by immunoblot **(Fig. 1C),** which recapitulates the haploinsufficiency of SHANK3 commonly described in patients.^15,46^ The 50 percent reduction of SHANK3 expression was also confirmed by immunohistochemistry with SHANK3 antibodies **(Fig. 1B-D)**.

### Removing *Shank3* from the NAc diminishes social preference

We first used the 3-chamber social interaction task to assess how conditional, NAc *Shank3* deletion and conventional *Shank3*^Δe4–22^ deletion alters social preference **(Fig. 2A)**.^47^ We found that WT mice showed a preference for the mouse chamber, but *Shank3*^Δe4–22^ mice did not show a preference to either chamber **(Fig. 2B)**. Similarly, WT mice preferred to be in close proximity to the mouse cup compared to the empty cup. *Shank3*^Δe4–22^ mice did not prefer either close proximity zone. **(Fig. 2C)**. We found no changes in distance traveled during the test between WT and *Shank3*^Δe4–22^ mice **(Fig. 2D)**. In the conditional deletion, *Shank3^fl/fl^ ^+CRE^* mice showed a significant preference for the empty chamber while *Shank3^fl/fl^ ^+eGFP^* mice preferred the mouse chamber **(Fig. 2E)**. Additionally, *Shank3^fl/fl^ ^+CRE^* mice spent significantly less time in the mouse chamber than *Shank3^fl/fl+eGFP^*controls **(Fig. 2E)**. *Shank3^fl/fl+eGFP^* mice also have a preference to be in close proximity with the mouse cup and spent significantly more time in close proximity of the mouse cup compared to *Shank3^fl/fl+CRE^*mice **(Fig. 2F**). *Shank3^fl/fl+CRE^* mice travel less distance than *Shank3^fl/fl+eGFP^* mice during the 3-chamber task **(Fig. 2G)**.

**Figure 2:**
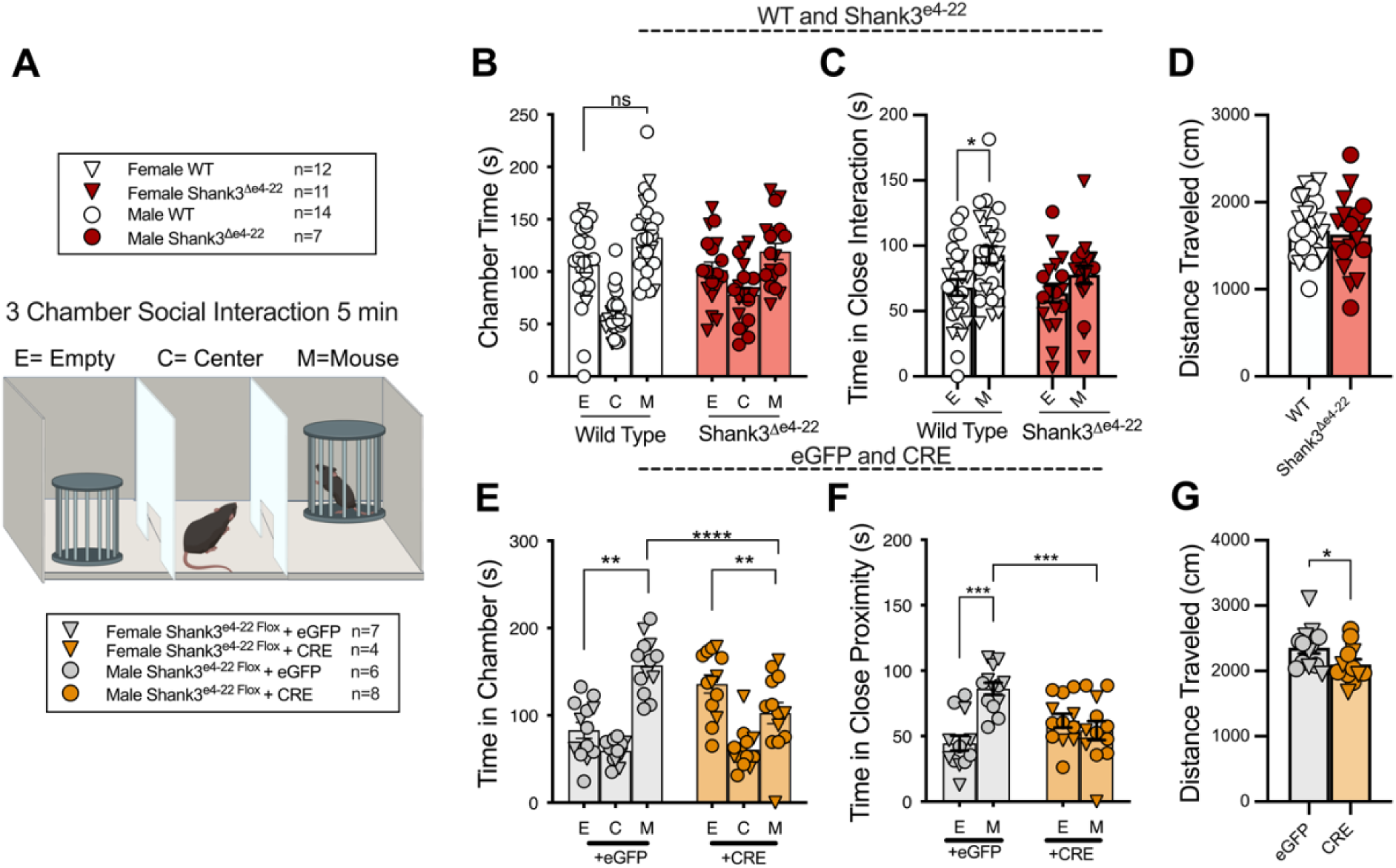
Conventional *Shank3^Δe^*^4–22^ Deletion and Conditional *Shank3*-NAc Deletion Eliminates Social Preference. **A)** Schematic of 3-chamber arena **B)** WT mice prefer the mouse chamber but not *Shank3^Δe^*^4–22^ mice (*\*P=0.012*). (Two-way ANOVA, Genotype: *F_(1,42)_=6.476; P=0.0147; n1=26, n2=18*) **C)** WT mice also spent significantly more time in close proximity (∼5cm perimeter) to the mouse-containing cup compared to empty, but not *Shank3^Δe^*^4–22^ mice (*\*P=0.116)* (Two-way ANOVA, Genotype: *F_(1,42)_= 2.586; P=0.1153; n1=26, n2=18*). **D)** There was no effect on the distance traveled between groups in the 3-chamber task (Unpaired t-test, Two-tailed, *P=0.7088*). **E)** *Shank3^fl/fl-eGFP^* mice showed a preference for the mouse chamber (\*\**P=0.0012*), whereas *Shank3^fl/fl+-CRE^* mice showed a preference for the empty chamber (*\*\*P=0.0078*). *Shank3^fl/fl+-CRE^* mice spend significantly less time in the mouse chamber than eGFP-injected mice (*\*\*\*P<0.0001*) (Two-way ANOVA, Chamber x Virus: *F_(1,23)_= 25.70; P<0.0001; n1=13, n2=12*). **F)** *Shank3^fl/fl-eGFP^* mice, but not *Shank3^fl/fl+CRE^* mice, spend significantly more time closely interacting with the mouse cup than the empty cup (\*\*\**P=0.0002*). *Shank3^fl/fl+eGFP^* mice spend more time in the mouse close interaction zone than *Shank3^fl/fl+CRE^* mice (*\*\*\*P=0.0005*) (Two-way ANOVA, Chamber x Virus: *F_(1,23)_= 15.28; P=0.0007; n1=13, n2=12*). **G)** *Shank3^fl/fl+CRE^* mice travel significantly less distance than *Shank3^fl/fl+eGFP^* mice. (Unpaired t-test, Two-tailed, *P=0.0435 n1=13, n2=12*). Complete statistical analyses are provided in the statistical analyses file associated with this manuscript.

Next, we extended our investigation to understand how *Shank3* deficiency impacts a naturalistic social environment using the juvenile social dyadic task (**Fig. 3A**). Mice were paired with sex-matched juvenile mice. *Shank3*^Δe4–22^ mice showed increased body contact time **(Fig. 3B)** compared to WTs. However, there was no difference between groups in the average distance between the test mouse and the juvenile target **(Fig. 3C)** or time spent in side-by-side facing the same direction **(Fig. 3D)**. *Shank3*^Δe4–22^ mice spent significantly more time grooming **(Fig. 3E)**. *Shank3^fl/fl+CRE^* mice were not different from *Shank3^fl/fl+eGFP^* controls in body contact time **(Fig. 3F)**, average distance from juvenile mouse (**Fig. 3G)**, time in side-by-side interaction **(Fig. 3H)**, or in grooming time **(Fig. 3I)**. Taken together, these data indicate that *Shank3* in the NAc does not underlie the time the animal spends in naturalistic social engagement.

**Figure 3:**
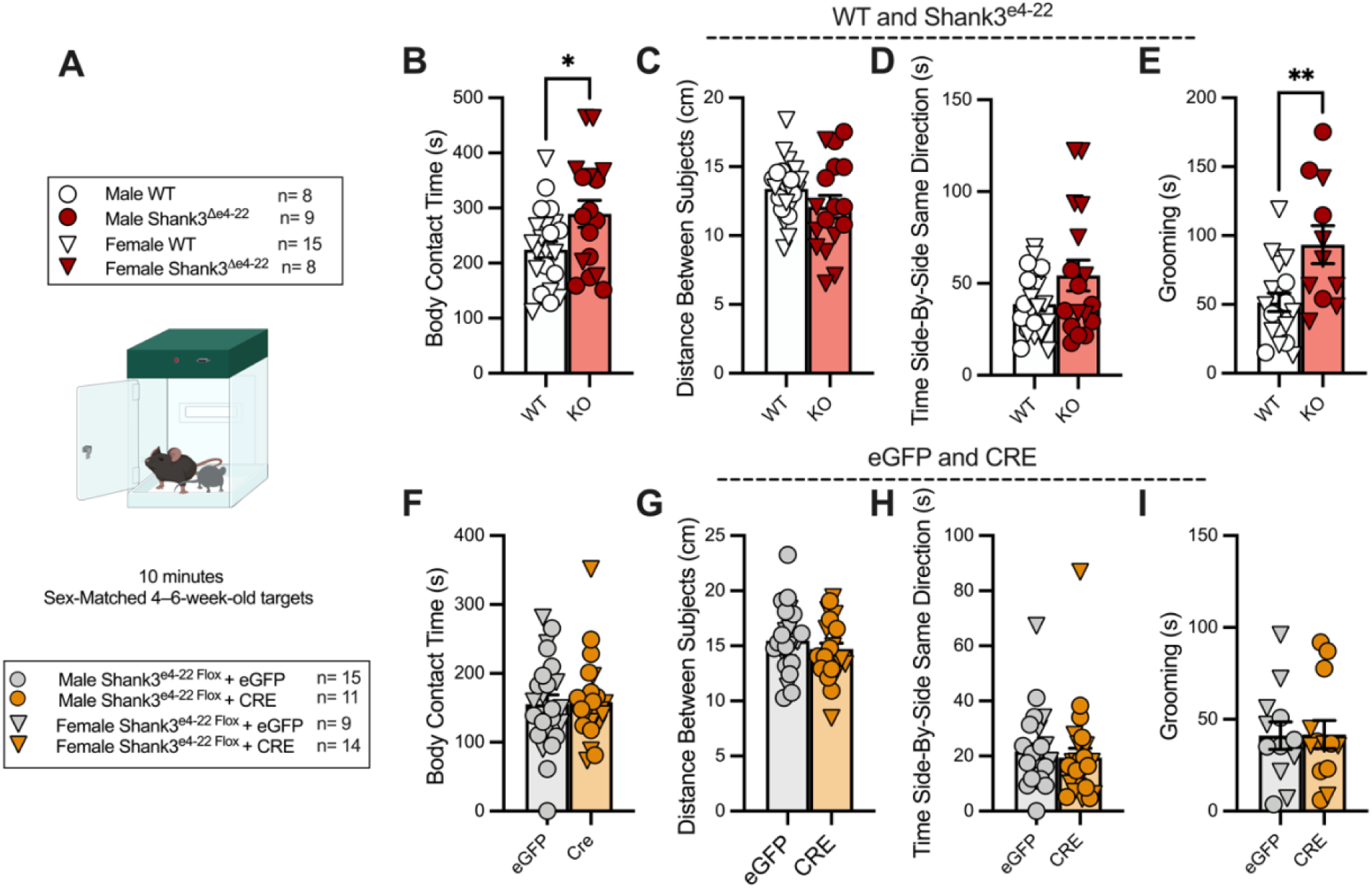
*Shank3^Δe^*^4–22^ Mice Show Increased Social Contact While *Shank3-*NAc Deletion Does Not Alter Dyadic Social behavior. **A)** Schema of experimental design of the social dyadic task. **B)** *Shank3^Δe^*^4–22^ mice showed increased time in body contact with a same-sex juvenile mate than WT controls (\**P=0.0225,* Unpaired t-test, Two-tailed, *n1= 23, n2= 17*). **C)** No difference between groups in the average distance between test mouse and juvenile target during testing (*P=*0.1735, Unpaired t-test with Welch’s correction, Two-tailed, *n1= 23, n2= 17*). **D)** No difference for the time in side-by-side facing the same direction between subjects during testing (*P=*0.0986, Unpaired t-test with Welch’s correction, Two-tailed, *n1= 23, n2= 17*) **E)** A random selection of *Shank3^Δe^*^4–22^ and WT mice were hand scored for grooming. *Shank3^Δe^*^4–22^ mice spend significantly more time grooming (\*\**P=0.0050,* Unpaired t-test, Two-tailed, *n1= 18, n2=11*). **F-I)** No differences between eGFP and CRE groups in time spent in body contact **(F)** (*P=*0.8594, Unpaired t-test, Two-tailed, *n1= 24, n2=25*), distance between subjects **(G)** (*P=*0.3537, Unpaired t-test, Two-tailed, *n1= 24, n2=25*), or time side-by-side same direction **(H)** (*P=*0.6177, Unpaired t-test, Two-tailed, *n1= 24, n2=25*). **I)** A random selection of grooming was hand scored for *Shank3^fl/fl-eGFP^* and *Shank3^fl/fl+CRE^* mice. No changes were detected in time grooming (*P=*0.9524, Unpaired t-test, Two-tailed, *n1= 12, n2=13*.) Complete statistical analyses are provided in the statistical analyses file associated with this manuscript.

### Removing *Shank3* from the NAc does not alter locomotion, grooming, or anxiety-like behavior

Previously, we showed that conventional *Shank3^Δe^*^4–22^ mice are hypoactive in the open field task, show robust hyper grooming, and increased anxiety-like behavior.^17^ We sought to rule out the possibility that changes in basic locomotive behaviors, grooming, or anxiety-like behaviors contribute to the loss of social preference seen in Cre-injected *Shank3^fl/fl^*mice by running a 10-minute open field trial **(Fig. 4A)**. We first recapitulated our previous findings and show that *Shank3*^Δe4–22^ mice travel less distance **(Fig. 4B)**, spend less time in the center **(Fig. 4C)**, enter the center fewer times **(Fig. 4D)**, and groom more **(Fig. 4E)**. However, in *Shank3^fl/fl^ ^+CRE^* mice, we found an increase in distance traveled compared to *Shank3^fl/fl+eGFP^***(Fig. 4F)**. We did not find any differences for the time spent in the center **(Fig. 4G)**, or entries to the center **(Fig. 4H)**. We also extended the open field to 30 minutes in these animals to look for changes in habituation over the longer trial **(Fig. S2F)**,^48^ as we previously reported hypolocomotion and decreased center time in *Shank3*^Δe4–22^ mice compared to WT during an extended open field trial.^17^ We did not see any differences between *Shank3^fl/fl+eGFP^ and ^+CRE^* mice in the 30-minute trial in the distance traveled **(Fig. S1G, I)**, time in the center **(Fig. S1H, J)**, or grooming **(Fig. S1 K)**. We also ran the Elevated Plus Maze (EPM) task **(Fig. S1A)** and found no differences between *Shank3^fl/fl^ ^+eGFP^ and ^+CRE^* mice in time spent in open arms, **(Fig. S1B)** closed arms, **(Fig. S1C)**, time in center **(Fig. S1D)**, or distance traveled **(Fig. S1E)**.

**Figure 4:**
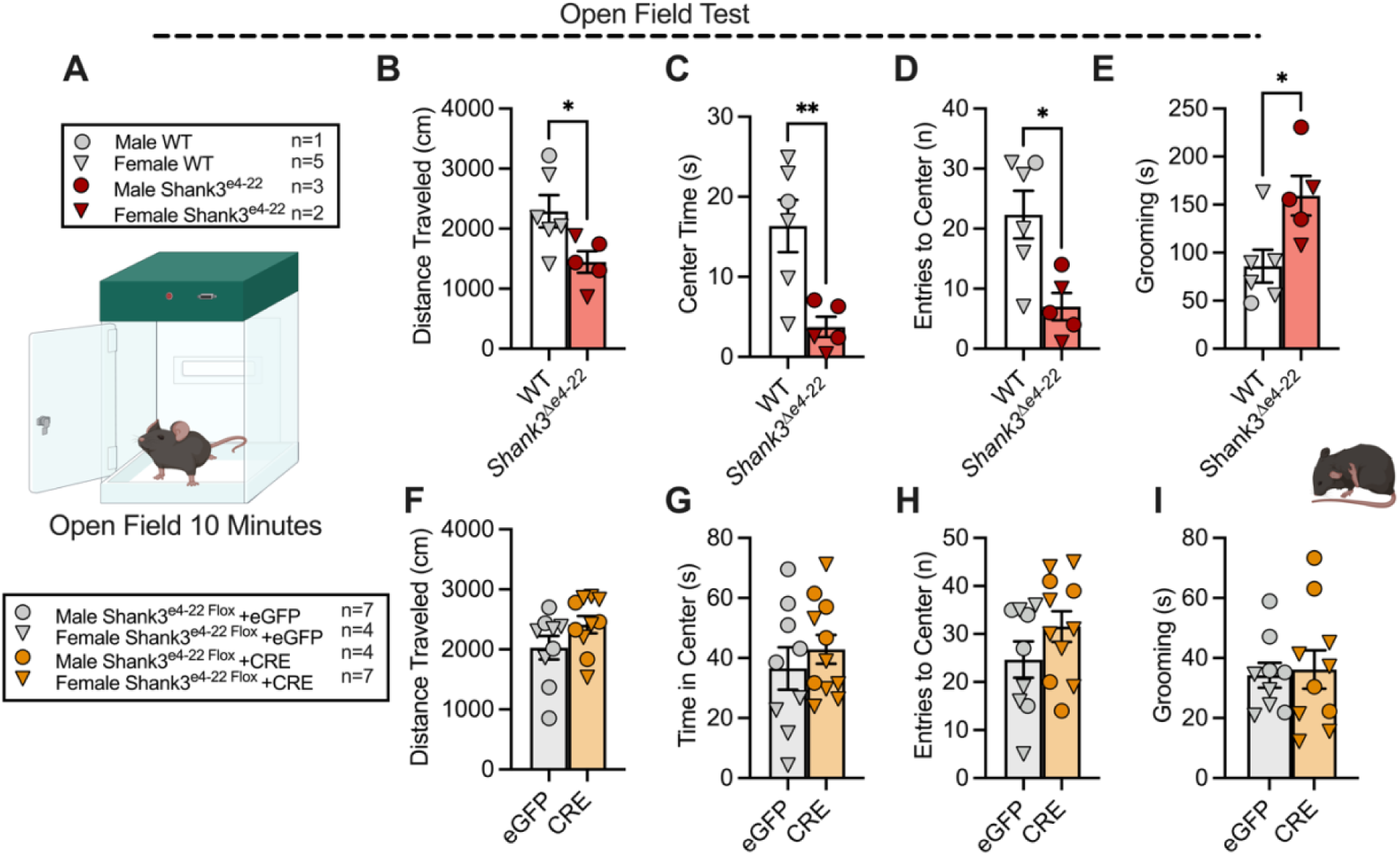
*Shank3^Δe^*^4–22^ Mice Show Reduced Locomotion and Increaed Anxiety-like Behavior, but *Shank3*-NAc Deletion Does Not Alter Anxiety-like Behaviors. **A)** Diagram of open field task **B-E) (B)** *Shank3^Δe^*^4–22^ mice traveled significantly less distance (*\*P=*0.0345, Unpaired t-test, Two-tailed, *n1= 6, n2=5*); **(C)** spent less time in the center (*\*\*P=0.0088,*Unpaired t-test, Two-tailed, *n1= 6, n2=5*); **(D)** had fewer entries to the center (\**P=0.0116*, Unpaired t-test, Two-tailed, *n1= 6, n2=5*); and **(E)** groom more (\**P=0.0215*, Unpaired t-test, Two-tailed, *n1= 6, n2=5*) compared to WTs. **F-H) (F)** There is no difference between *Shank3^fl/fl+CRE^* and *Shank3^fl/fl+eGFP^* mice in distance traveled (*P=0.1286,* Unpaired t-test, Two-tailed, *n1= 9, n2=10*), **G)** time spent in the center (*P=0.4575,* Unpaired t-test, Two-tailed, *n1= 9, n2=10*) **H)** entries to the center (*P=0.1789,* Unpaired t-test, Two-tailed, *n1= 9, n2=10*) or **I)** grooming (*P=0.8077*, Unpaired t-test, Two-tailed, *n1= 9, n2=10*). Complete statistical analyses are provided in the statistical analyses file associated with this manuscript.

### Removing *Shank3* from the NAc decreases social reward-seeking but not food reward-seeking

The NAc is a well-established modulator of motivation,^33^ and we have shown that *Shank3^Δe^*^4–22^ mice have depleted motivation during the rewarded lever pressing tasks.^17^ Therefore, we next tested *Shank3^fl/fl^*mice in a rewarded lever press task (**Fig. 5A-B)**. *Shank3^fl/fl+CRE^* mice showed no differences in the number of lever presses across 6 training days (**Fig. 5C)** or the test day (**Fig. 5C,D)** compared to *Shank3^fl/fl+eGFP^* mice. To assess any differences in motivation, we extended the lever press task to include a breakpoint task on day 8.^49^ We found no differences between *Shank3^fl/fl^*^+CRE^ and *Shank3^fl/fl+eGFP^*mice **(Fig. 5E)**.

**Fig 5:**
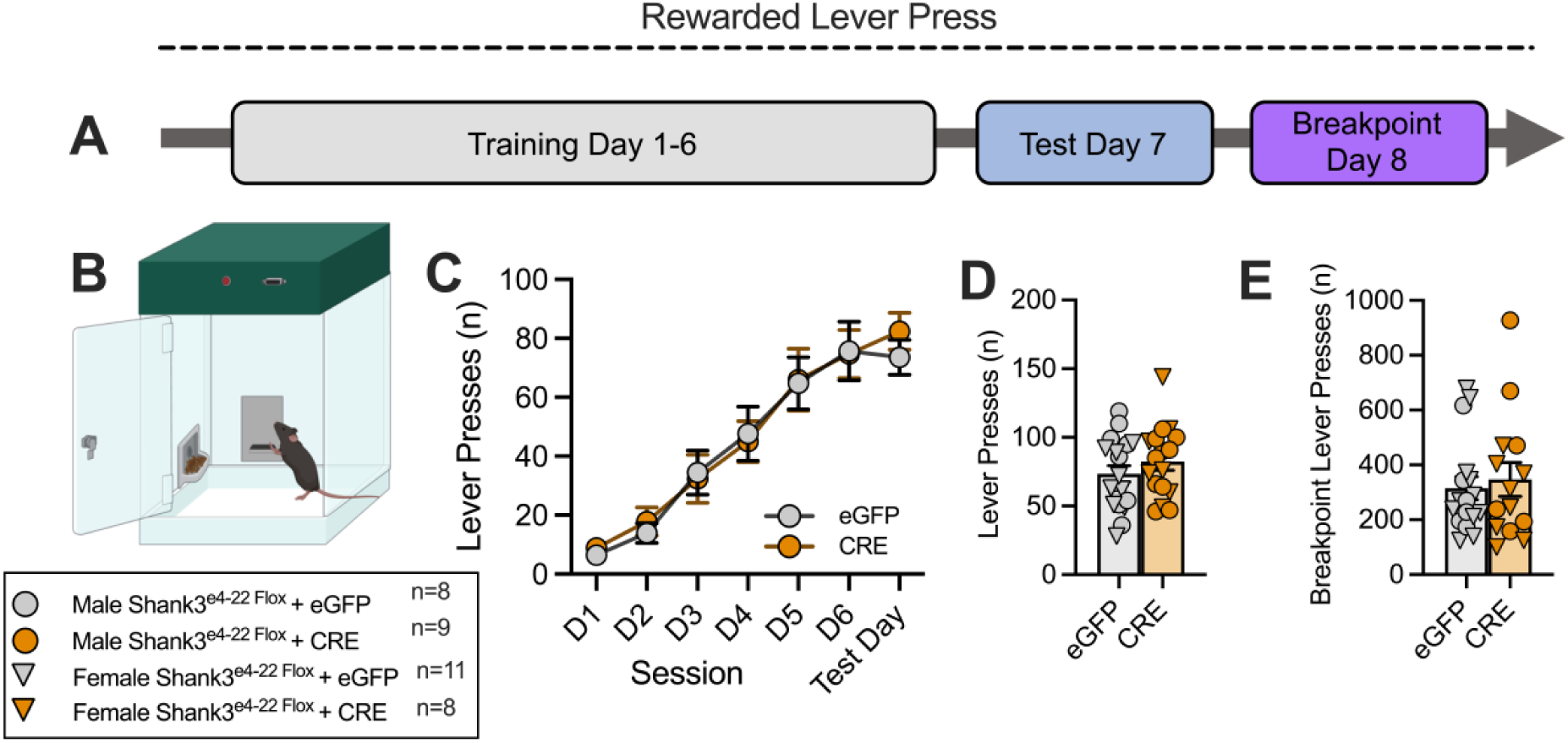
Removing *Shank3* from the NAc does not affect palatable food reward-seeking behavior. **A)** Timeline of experiment **B)** Schema of lever press apparatus. **C)** Lever pressing counts do not differ between *Shank3^fl/fl+CRE^* and *Shank3^fl/fl+eGFP^* mice across the 6 days of training or test day of the experiment (Two-way ANOVA, Day x Virus: *F_(6,204)_=0.2230; P=0.9690; n1=19, n2=17*). **D-E) D)** There are no differences in the number of lever presses of animals between *Shank3^fl/fl+CRE^* and *Shank3^fl/fl+eGFP^* mice on test day (Unpaired t-test, Two-tailed; *P=0.315; n1=19, n2=17*) or **E)** during breakpoint on day 8 (Unpaired t-test, Two-tailed; *P=0.6689; n1=17, n2=14*). Complete statistical analyses are provided in the statistical analyses file associated with this manuscript.

Given that conditional removal of *Shank3* from the NAc and conventional *Shank3^Δe^*^4–22^ mutation decreases social investigation in the 3-chamber task, we predicted that *Shank3^Δe^*^4–22^ and *Shank3^fl/fl+ CRE^* mice are less rewarded by social interaction and, therefore, show less motivation to enter the mouse chamber. To test this hypothesis, we used a social condition place preference (sCPP) assay to measure social motivation (**Fig. 6A)**.^50^ WT mice show a significant increase in time spent in the social chamber compared to empty in the post-test, but not the pre-test **(Fig. 6B)**. Interestingly, we also found *Shank3^Δe^*^4–22^ mice spend more time in the social-paired chamber on post-test day **(Fig. 6C)**. *Shank3^fl/fl+eGFP^*mice showed a trend toward spending more time in the social-paired chamber on post-test day **(Fig. 6D)**. Notably, *Shank3^fl/fl+CRE^*mice showed a preference for the empty-paired chamber on post-test day, revealing an opposite result from all other groups. **(Fig. 6E)**. When we compared the time in the social-paired chamber on post-test day, we found *Shank3^fl/fl+CRE^* mice spend significantly less time in the social-paired chamber compared to *Shank3^fl/fl+eGFP^* mice **(Fig. 6F)**. Similarly, when we calculated the CPP score, *Shank3^fl/fl+CRE^* mice had significantly lower scores than *Shank3^fl/fl^ ^+eGFP^* and WT mice **(Fig. 6G)**. Moreover, we reported a significant difference between the groups in standard deviation, as calculated by the Brown-Forsythe test **(Fig. 6G)**. *Shank3^Δe^*^4–22^ mice are hypoactive on the pre-test day but also show a significant decrease in the distance traveled on the pre-test day compared to the post-test day **(Fig. 6H)**, which we did not find in any other group **(Fig. 6I)**. We also measured body contact duration in males and social sniffing in females of our four test groups across the 4 days of conditioning, and we did not find any significant differences across all groups **(S3A-F)**.

**Figure 6:**
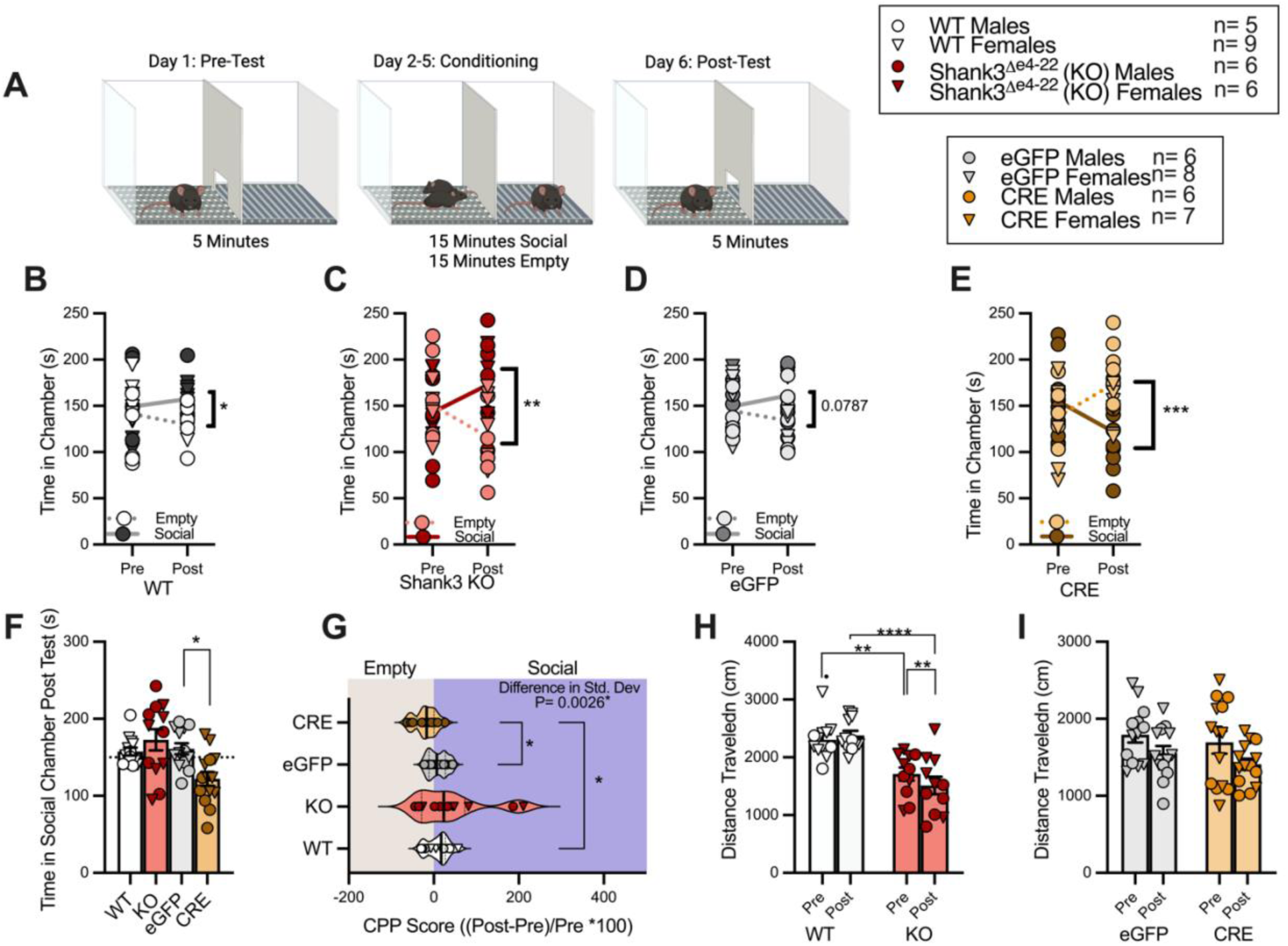
Removing *Shank3* from the NAc Causes Social Conditioned Place Aversion. **A)** Social Conditioned Place Preference (SCPP) Experimental Design **B-C) (B)** WT (*\*P=0.0106.* Mixed-effects analysis, Chamber: *F_(1,48)_=0.04345; P=0.0196; n=13*). and **(C)** *Shank3^Δe^*^4–22^ mice (Post Empty vs. Social *\*\*P=0.0018,* Mixed-effects analysis, Time x Chamber: *F_(1,44)_=7.155; P=0.0105; n=12*) spend significantly more time in the social-paired chamber in the post-test. **D)** There was a trend toward increased time in the social-paired chamber for *Shank3^fl/fl+eGFP^* mice. (Post Empty vs Social: *P=0.0787;* Mixed-effects analysis, Time x Chamber: *F_(1,12)_=3.367; P=0.0914; n=13*) **E)** *Shank3^fl/fl+CRE^* mice spend significantly more time in the empty-paired chamber that social-paired during the post-test day (Post Empty vs. Social: *\*\*\*P=0.0006;* Mixed-effects analysis, Time x Chamber: *F_(1,13)_=17.06; P=0.0012; n=14*) **F)** *Shank3^fl/fl+CRE^* mice spend significantly less time in the social-paired chamber on the post-test day than *Shank3^fl/fl+eGFP^* mice (*\*P=0.0141;* Brown-Forsythe ANOVA test, *F_(3, 30.94)_=5.696; P=0.0032; n1=13, n2= 12, n3=13, n4=14*). **G)** *Shank3^fl/fl+CRE^* mice have a significantly lower CPP score than WTs (*\*P=0.0479*) and eGFP-injected mice (*\*P=0.0359;* Brown-Forsythe ANOVA test, *F_(3, 17.02)_=3.258; P=0.0474; n1=13, n2= 12, n3=13, n4=14*). There was also a significant difference in the standard deviation of groups in this task (Brown-Forstythe test *P=0.0026)* **H)** Distance traveled is significantly lower in *Shank3^Δe^*^4–22^ mice compared to WTs during the pre-test (*\*\*P=0.0051*) and post-test (*\*\*\*\*P<0.0001). Shank3^Δe^*^4–22^ mice also travel less distance in the post-test compared to pretest (*P=0.0074*) (Mixed-effects analysis, Time x Chamber: *F_(1,22)_=6.738; P=0.0165; n1=12, n2=12*). **I)** There are no changes in distance traveled in the pretest to posttest analysis or between *Shank3^fl/fl+eGFP^* and *Shank3^fl/fl+CRE^* mice (Mixed-effects analysis, Time x Virus: *F_(1,25)_=0.04529; P=0.8332; n1=13, n2=14*). Complete statistical analyses are provided in the statistical analyses file associated with this manuscript.

## DISCUSSION

In the present study, we showed that removing *Shank3* from the NAc selectively eliminates social motivation but does not affect naturalistic social behaviors, motivation for palatable food, or anxiety-like behaviors, all of which are present in the conventional *Shank3^Δe^*^4–22^ mouse model. Interestingly, we also report that *Shank3^Δe^*^4–22^ mice show decreased social preference but increased social motivation. We conclude that *Shank3* in the NAc is an important regulator for only social motivation. These data give a novel, mechanistic understanding of how social motivation is impaired in a genetically validated ASD model.

We initially anticipated that disruption of *Shank3* expression in the NAc would alter all motivation and social behaviors and recapitulate the blunted social and reward-seeking phenotypes of *Shank3^Δe^*^4–22^ mice. However, we showed that removing *Shank3* from the NAc only disrupts social motivation. We then predict that *Shank3* in NAc-connected regions regulates food motivation and naturalistic, reciprocal social interaction. The ventral tegmental area (VTA) is a prime site for further investigation as it shares a reciprocal circuitry with the NAc and is a well-established modulator of motivation. Knocking down SHANK3 in the VTA also disrupts social preference.^51^ However, *Shank3*-VTA deletion mice also lack sucrose preference, indicating a non-specific loss of reward-seeking behavior.^51^ Taken together with the present study, SHANK3 is likely necessary for social motivation at both ends of the NAc-VTA circuit, but removing SHANK3 has a different effect on neural activity and encoding of behavior on each side of the circuit. Beyond social motivation, removing *Shank3* from the vCA1 of the hippocampus disrupts social memory in mice, and removing *Shank3^Δ^*^14–16^ from the anterior cingulate cortex (ACC) disrupts social preference in the 3-chamber assay.^52,53^ The vCA1 of the hippocampus and the ACC project to the NAc and regulate social behaviors^54,55^ There may be a convergent mechanism where the loss of *Shank3* at one loci of social brain networks is sufficient to change social behavior due to its influence across the network. Further investigation of the effect of *Shank3* NAc deletion on neural activity across brain-wide networks, particularly those involved in social motivation, will provide a holistic mechanistic understanding of social motivation and social behaviors.

It was unanticipated that *Shank3^Δe^*^4–22^ mice would show decreases in social preference in the 3-chamber task but increases in time in the social-paired chamber in the sCPP task. Yet, when we consider the increased anxiety-like behaviors in *Shank3^Δe^*^4–22^ mice reported here and in our previous study^17^, we predict that the heightened stress levels confound the results seen in the sCPP as a negative affective state is shown to influence social motivation in mice and patients with ASD.^56,57^ On test day, we see significant increases in data variability in *Shank3^Δe^*^4–22^ mice. This is reminiscent of past reports showing that stressing mice before running a CPP assay causes significant increases in side-preference variability and decreases in the distance traveled.^58^ This report concludes that stress drives mice to randomly choose a side, independent of training, and remain there for most of the task. We predict that the chronic, novel, free social interaction may be stress and anxiety-inducing for *Shank3^Δe^*^4–22^ mice and drive this phenotype.^54^ *Shank3B^-/-^* (*Shank3^Δ^*^14–16^) mice also show increased anxiety-like behavior but decreased social reward, as shown in an operant task.^18,54,59,60^ Future studies should examine how *Shank3^Δe^*^4–22^ mice perform in a social-rewarded operant task, which eliminates free interaction, to determine if social dyadic behavior confounds social motivation. Furthermore, we show that *Shank3^fl/fl+CRE^* mice develop a conditioned place aversion to the social paired chamber and do not show any changes in baseline anxiety-like behaviors. Without baseline changes in affective states in our conditional deletion confounding social motivation, we posit that *Shank3* in the NAc directly underlies social motivation.

This also led us to question how neural circuitry underlies this communication between anxiety-like states and social motivation. Others have shown that removing *Shank3* only from the Bed Nucleus of the Striata Terminalis (BNST) replicates the anxiety-like behaviors of *Shank3^Δe^*^4–22^ mice.^61^ The BNST-NAc circuit regulates stress adaptations and social behaviors in a higher-stress state and, therefore, could regulate the influence of stress on social motivation.^62^ Future experiments should investigate beyond one brain region to understand how the social brain integrates the affective state with social motivation.^63^

We establish a new *Shank3^fl/fl^* mouse model with two loxP sites flanking exons 4 and 22 that significantly improves recombination efficiency with a predictable pattern related to diverse *Shank3* isoforms. Phelan-McDermid Syndrome (P-MS) or 22q13.3 deletion syndrome, is defined by a loss of the distal long arm of one chromosome 22 which includes the gene SHANK3 and the majority of patients are diagnosed with autism.^64^ Our findings show that we successfully knockdown SHANK3 protein expression to 50% of WTs, which recapitulates the haploinsufficiency of SHANK3 found in patients.^15,46^ This model thus provides a novel avenue for exploring conditional SHANK3 deficiency that replicates the genetic abnormalities found in patients. However, findings from our recent study of WT, *Shank3^Δe^*^4–9^, and *Shank3^Δe^*^4–22^ mice using a targeted RNA capture and long-read sequencing method presented a more complex structure for WT and mutant *Shank3* transcripts^65^. We unexpectedly find fusion transcripts of *Shank3* coding exons and the downstream protein coding gene *Acr* in WT mice. *Shank3^Δe^*^4–22^ deletion also results in highly expressed fusion transcripts between non-deleted exons of *Shank3* and the coding exons of *Acr* that are not expressed in WTs. The function of these fusion transcripts remains to be defined in WT and mutant mice *Shank3^Δe^*^4–22^ deletion disrupts the largest portion of the coding region within the previously characterized *Shank3* transcripts^17,66^. *Shank3^Δe^*^4–22^ deletion recapitulates the loss of function of known SHANK3 protein and has molecular validity and recapitulates the *SHANK3* mutations found in ASD patients.^14,15^ Therefore, *Shank3*^Δe4–22^ and our new *Shank3^fl/fl^* mice are appropriate tools for investigating the pathomechanisms of *SHANK3*-associated ASD. However, it remains to be investigated whether *Shank3*^Δe4–22^ deletion recapitulates the *SHANK3* deficiency in P-MS patients because of the fusion transcripts.

Here, we conditionally deleted *Shank3* in the NAc of adult mice. This approach gave us the advantage of high-precision targeting in the NAc to establish our initial findings. Future study is warranted to determine how loss of *Shank3* in the NAc during early development affects social motivation, as RNA silencing of *Shank3* in the NAc eliminates social preference, but only when done at P6.^27^ Further investigation of the knockdown of SHANK3 in the NAc at additional developmental time windows is needed to provide a comprehensive understanding of how NAc-SHANK3 coordinates social motivation.

Overall, these data establish a novel and specific role of *Shank3* in the NAc on social motivation and support a growing body of evidence that social motivational mechanisms may underlie autism phenotypes. The social motivation theory of autism is a classic theory for understanding social deficits in autistic individuals,^63^ but recent publications argue that increased anxiety and overinterpretation of human behavior could be confounding our understanding of social motivation in autism.^57,67,68^ Our data establish a prospective mechanism and model for how co-occurring anxiety-like phenotypes can shift social motivation and provide an avenue to investigate these hypotheses.

## METHODS

### Animals

Mice were group housed by sex on a 12-hour light/ 12-hour dark cycle with lights on at 0700 hours. Food and water were available ad libitum, except during rewarded lever press experiments (see experimental details). All experiments were conducted during the light phase. Animal husbandry and behavioral testing and animal husbandry protocols were approved by the Yale Animal Resources Center and in accordance with the Institutional Animal Care and Use Committee.

*Shank3^fl/fl^* mouse line was engineered via CRISPR/Cas genome editing. In our previous *Shank3 e4-22^flox/flox^* mouse line^45^, three loxP sites were located at between intron 3, 9, and downstream of exon 22 of *Shank3* genes (GenBank:NM_033517). A sgRNA was designed to target the second loxP site in *Shank3 e4-22^flox/flox^* mouse line. The validation of sgRNA was done *in vitro*. After pronuclear injection of the Cas9 and sgRNA vector, PCR and genomic sequencing was done to confirm the deletion of the second loxP sequence and the presence of the first and third loxP sequence in the founder mouse. The founder mouse was then bred with a C57/BL6 mouse to generate heterozygous offspring. The litters were genotyped using PCR and the pup containing two loxP sites was bred with C57/BL6 mice for more than 5 generations to eliminate any potential off-target events. No unusual phenotypes were observed in the animals. After ∼10 generations of breeding, we used homozygotes breeding (*Shank3^fl/fl^ X Shank3^fl/fl^*) pairs to maintain the mouse line and obtain the experimental mice.

### Genotyping and PCR confirmation of loss of *Shank3*

Primers FLP-Neo-F: (5’-gggaggattgggaagacaat-3’), SH-AFLP-R (5’-ggctatgttcatgggatcttgt-3’), and SH-BFLP-F: (5’-ttgccgaggtaatcaagacc-3’) were used to amplify WT (390bp) and *Shank3^fl/fl^* alleles (211bp). To identify recombination of the loxP sites (Δe4-22), primers SH3-MTF (5’-ttgcatctgggacctactcc-3’), SH3-MTR (5’-aaagcactgactcctctcttgg-3’), SH3-WTF (5’ gtgccacgatcttcctctaaac-3’), SH3-WTR (5’ agctggagcgagataagtatgc-3’) were used to amplify a 200bp product when the floxed allele was intact and a 600bp product when recombination had occurred with a 30s denaturing step at 94°C, 30 s annealing step at 59°C, and a 45 s extension step at 72°C for 40 cycles.

### Viruses

*Shank3^fl/fl^* mice were injected with 300nL pAAV.CMV.HI.eGFP-Cre.WPRE.SV40 or 300nL AAV5.hSyn.eGFP.WPRE.bGH (Addgene). pAAV.CMV.HI.eGFP-Cre.WPRE.SV40 (Addgene viral prep # 105545-AAV5; http://n2t.net/addgene:105545; RRID:Addgene_105545) and pAAV.CMV.PI.EGFP.WPRE.bGH (Addgene viral prep # 105530-AAV5; http://n2t.net/addgene:105530; RRID:Addgene_105530). Viral preps were gifts from James M. Wilson.

### Stereotaxic Surgery

Male and female *Shank3^fl/fl^* mice underwent bilateral stereotaxic surgery at 6-12 weeks of age. Animals were initially anaesthetized at 5% isoflurane and administered Ethiqa XR 1.3mg/mL at 0.6mg/kg (S.Q., Covetrus) for preemptive analgesia. Animals were then moved to the stereotaxic frame (RWD) and anesthesia was maintained on 1% isoflurane, 1% oxygen. Following a craniotomy, viruses were loaded into a NanoFil 10uL syringe equipped with a 33-gauge blunt needle tip (World Precision Instruments). The needle was moved into the NAc (anterior/posterior [AP]: –1.50mm, medial/ lateral [ML]: +/− 0.75mm, dorsal/ ventral [DV]: 4.50mm), paused for 1 minute, then moved to DV 4.45mm and the virus was infused into the NAc at a rate of 100nL/ minute with an automated infuser (Harvard Instruments). Following infusion of virus, the syringe remained in place for 10 minutes and was then removed from the brain. Following bilateral injection, the incision site was sutured, and mice were given 1mL of saline S.Q. and 5mg/kg Carprofen I.P. Mice received 5mg/kg injections of carprofen every 24 hours for 72-hour recovery monitoring. Mice recovered for 3-4 weeks before any experiments.

### Immunoblot

<u>Tissue</u>: 4 weeks after viral infusion, NAc brain tissue was collected. Mice were anesthetized using isoflurane in a bell jar then rapidly decapitated and the brain was extracted on ice. The brain was placed in a cold brain block (Harvard Instruments) and sections containing the NAc were isolated between razor blades. NAc sections were frozen on dry ice and the NAc was isolated using a 2mm biopsy punch (Integra Miltex) and stored in an Epindorf tube in –80°C.

<u>Preparation of crude synaptic proteins:</u> The NAc tissue was homogenized in Syn-PER Synaptic Protein Extraction Reagent solution (Thermo Scientific) with protease and phosphatase inhibitors. Homogenate was centrifuged at 1200 x *g* for 10 minutes at 4°C. The supernatant was separated and centrifuged at 15000 x *g* at 4°C for 20 minutes. This synaptosome pellet (P2) was suspended in Syn-PER Reagent, and the cytosolic fraction (S2) was saved.

<u>Quantitative immunoblot analysis:</u> immunoblot analysis was conducted following previous reports.^17,45^ Equal protein amounts from the synaptosome and cytosolic fraction were separated by SDS-PAGE and transferred to PVDF membranes. The membranes were blocked for 1 hour in 0.02M Tris-Buffered Saline (TBS) with 0.1% Tween-20 (TBST) and 5% non-fat milk solution then incubated overnight at 4°C in primary antibody [Shank3, rabbit (Courtesy of Yong Q. Zhang lab at Hubei University, 1:1000) or β actin (Santa Cruz sc-1615 1:5000)]. The blots were then washed with TBST 3 times for 10 minutes each and incubated with HRP-conjugated secondary antibody [Anti rabbit HRP (Santa Cruz, 1:3000) or Anti mouse HRP (Santa Cruz, 1:5000)] (for 1 hour at room temperature. The membrane was then washed 3 times for 10 minutes in TBST. The membrane was incubated with ECL reagent for 1 minute and imaged using a Chemidoc (BioRad). Band sizes were analyzed using ImageJ and normalized to β-actin loading control.

### Histology and Imaging

<u>Tissue:</u> After completion of behavior experiments brain tissue was collected. Mice were anesthetized using isoflurane in a bell jar and transcardially perfused with phosphate buffered saline (PBS, 10mL) followed by 4% paraformaldehyde in 0.1 phosphate buffer (PFA, 15-20mL). Brains were harvested and stored overnight in 4% PFA and transferred the following day to a 30% sucrose solution for at least 48 hours. Brains were cut at 80uM using a Leica CM3050 S cryostat or a Leica SM2000R microtome and slices were stored in an antifreeze, ethylene-glycol solution and stored at –20° until analysis.

<u>Viral Validation Experiments:</u> Tissue was transferred to cell strainers in a bath of fresh PBS for 10 minutes 4 times. Slices were then mounted onto charged slides and mounted in Dapi-Fluromount-G (SouthernBiotech) and sealed with a coverslip and clear nail polish. Images were collected at 10x objective on a Zeiss 980 and manually reviewed for viral expression by a blind experimenter.

<u>Immunohistochemistry:</u> tissue was transferred to cell strainers and washed in a fresh bath of PBS for 10 minutes 4 times. Next, slices were transferred to a PBS + 0.3% Triton-X 100 solution and washed for 60 minutes. Slices were then washed in a blocking buffer (0.3% Triton-X 100 and 10% Blocking One (Nacalai USA, Inc.) in PBS filtered with 0.22uM filtering unit) for 60 minutes.

Slices were then transferred to primary antibody solution overnight at room temperature. SHANK3 (Cell Signaling Technology #64555) solution was made at 1:1000 concentration in blocking buffer. The next day, the slices were washed 4 times in fresh PBS for 10 minutes each and then were incubated in Alexa 647 anti-rabbit (Fisher Scientific) at 1:1000 for 3 hours at room temperature. Slices were again washed 4 times for 10 minutes in fresh PBS then incubated in DAPI (1:1000, VWR) for 5 minutes and washed another 4 times for 10 minutes each in fresh PBS. Finally, tissue was mounted onto charged slides using Fluoromount-G (EMS Acquisition) and cover slips were sealed with clear nail polish.

<u>Imaging:</u> Images of slices were taken using a Zeiss 980 using 10x, 20x and 63x objective. Slices were compiled into a tiled image on one plane. For Shank3 expression analysis, images were first deidentified so the experimenter could remain unbiased, and the NAc brain region was identified using a Mouse Brain Atlas (Paxinos and Franklin). We then calculated corrected total cell fluorescence using ImageJ (corrected total cell fluorescence= Integrated Density – (Area * Mean Fluorescence of background)). All measurements were standardized as a percent change from total average eGFP readings. For each mouse, we averaged the corrected total cell fluorescence between the two NAc hemispheres to get the final result.

### Behavior Testing

All behavior experiments were recorded and analyzed using Noldus EthoVision XT (Leesburg, VA USA). All test and target mice were handled and tail marked for at least 3 days prior to testing. On the day of testing, animals were acclimated to the testing room for at least 1 hour prior to experiments. All equipment was cleaned prior to and in between trials using 70% etoh spray.

Cohorts:

1) <u>3-chamber social investigation test.</u> We used a 3-chamber arena comprised of a 420mm x 565mm x 358mm box with 185mm x 420mm equal chambers made of 5mm thick, clear Plexiglas with two doors (120mm wide, 5mm thick, 358mm tall) that connected chambers to the middle (Yale Machine Shop). The arena was outfitted with two empty wire pencil cups (Organize-it) that were flipped upside-down in the top corners of the arena with a 5cm path around each cup. Full water bottles were placed on top of the cups to secure them in place. Test mice were placed in the center chamber for 5 minutes with the doors closed to habituate to the arena. Next an age and sex-matched target mouse was placed underneath the cup in one of the external chambers. The doors were opened, and the test mouse was given 5 minutes to explore the arena. The location of the target mouse alternated for each subsequent trial. Target mice were acclimated to the pencil cups for 3 days prior to study. Time spent in each chamber, time spent within 5cm of each cup (called close interaction) and distance traveled was recorded. Experiment was conducted in 80-130 lux.
1) <u>Juvenile Social Dyadic Task:</u> Test mice and sex-matched juvenile (<5 week-old mice) were placed in the Noldus Phenotyper (Noldus) externally lined with white paper to obfuscate clear plexiglass walls. Mice were permitted to freely interact for 10 minutes. In addition to analysys in Ethovision, videos were analyzed post-hoc by a blinded coder using Behavioral Observation Research Interactive Software (BORIS)^69^ for fleeing or withdrawing from target, sniffing anogenital region, sniffing nose to nose, self-grooming, following target, and passively social as previously described.^23^
2) <u>Social Conditioned Place Preference:</u> The social conditioned place preference (sCPP) assay was modified from previous studies^50^. All studies were completed in red light (Lux <5). Animals were placed into a white plexiglass box (30cm x 30cm) fitted with two clear plexiglass inserts that created a 2-chamber arena. One chamber was lined with black and white stripped paper between the white plexiglass and clear liner and a metal wire cloth floor (McMaster, 85385T937) was placed on the floor. The second chamber was lined with black polka dot paper and a perforated steel floor with a staggered pattern (McMaster, 9358T221). The two chambers were connected by a clear Plexiglass door. On day 1, the door was removed, and mice were permitted to freely explore both chambers for 20 minutes. After each test the floors were washed with Alconox soap and water and dried with paper towels. The second day, in the first session (between 0900 and 1400 mice were placed into a randomly assigned side of the chamber with a novel, sex-matched juvenile target for 15 minutes (social-paired side) and kept on that side for the duration of the session with the clear door inserted. In the second daily session, from 1400 to 1900, mice were placed in the opposite side of the same box for 15 minutes alone (empty-paired side). On the third day mice were trained on empty-paired side for 15 minutes in the first session and social paired side in the second session. This alternating pattern of social and empty-paired training first continued for days 4 and 5. A new target mouse was used daily for training. Finally, on day 6, mice were placed in the chamber alone with the door removed and permitted to freely explore the 2-chamber arena for 20 minutes. For analysis, only the first 5 minutes of testing was used.
3) <u>Rewarded Lever Press:</u> This paradigm was adapted from previous reports.^17^ Mice were weighed 3 days prior to testing and then the food hopper was removed from cages for the duration of the task. Mice were given approximately 1g of food a day per mouse in the cage and weighed daily. Mice continued a restricted diet for the duration of the task and adjusted food availability to maintain 85-90% of starting body weight. On the day before testing, 5-10 20mg Dustless Precision Pellets (VWR) per mouse were placed into the home cage of animals to reduce neophobia. For testing, we used Noldus Phenotypers (Noldus) fitted with 1 wall that contained a lever, and 1 wall that contained a pellet receptacle attached to an automated dispenser filled with reward pellets. The lever was programmed to release 1 pellet for 1 lever press for the first 7 days of testing. On day 1, the lever was baited with 2 pellets and the pellet receptacle was baited with ∼3 pellets. Mice were given a daily session of 60 minutes for 7 days to freely explore and interact with the lever. On day 8, we ran a break point task in which the lever was programmed on a progressive ratio 5 schedule(e.g. 1 press per pellet then 5, then 10, etc). Number of lever presses were recorded daily.
4) <u>Elevated Plus Maze:</u> Mice were placed in the center of the elevated plus maze (San Diego Instruments) and permitted to freely explore the open arms (lux∼200) and closed arms (lux∼50) for 300 seconds. Time spent in open arms, closed arms, center, and total distance traveled was quantified.
5) <u>Open Field:</u> Mice were placed in the center of the Noldus Phenotyper fitted with 4 clear plexiglass walls and permitted to explore the chamber for 10 or 30 minutes freely. Total distance traveled, time in center (central 5cm square), and grooming (automatically detected by Ethovision XT) was quantified.

### Exclusion criteria

For all viral and implant studies, animals were excluded based on *a priori* standards. The injection site of all viral injections was identified by the presence of eGFP fluorescent marker. Animals were excluded from all data sets if the viral expression was not bilaterally expressed in the targeted region. Additionally, data were analyzed using Grubbs’ outlier test (alpha = 0.05) and any identified outliers were removed from the experiment.

### Statistics

Data are represented as mean ± SEM, with individual plot points overlaid on a mean bar graph. Annotation of * p<0.05, **p<0.01, and *** p<0.001 are used in figures. Statistical analysis was conducted using Prism 10 (GraphPad). Statistical tests and parameters are indicated in figure legends. Significance was set at α = 0.05, and a *P* value less than 0.05 was considered significant. Detailed statistic results and tests for all results can be found in the Supplementary Data.

## Supporting information

Statistical Analyses File

## ACKNOWLEDGEMENTS

**Figures were created in BioRender. Folkes, O. (2025) https://BioRender.com/p55k060**

