## Supplemental Data 1 for "Deficiency of *Shank3* in the Nucleus Accumbens Reveals a Loss of Social-Specific Motivation"

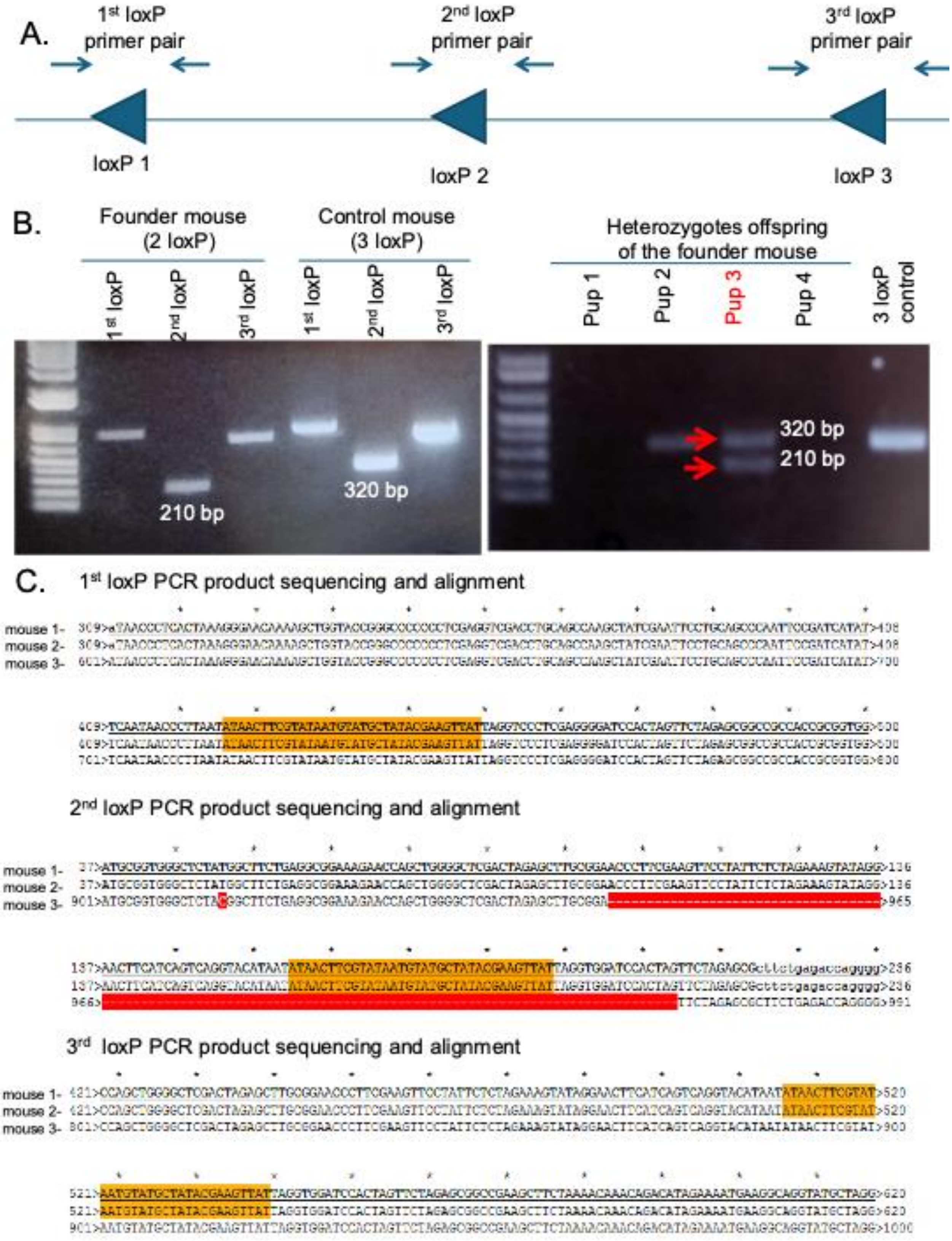
A.
1st loxP primer pair
2nd loxP primer pair
3rd loxP primer pair
loxP 1
loxP 2
loxP 3
B.
Founder mouse (2 loxP)
Control mouse (3 loxP)
Heterozygotes offspring of the founder mouse
1st loxP
2nd loxP
3rd loxP
Pup 1
Pup 2
Pup 3
Pup 4
3 loxP control
210 bp
320 bp
C.
1st loxP PCR product sequencing and alignment
mouse 1-
mouse 2-
mouse 3-
2nd loxP PCR product sequencing and alignment
3rd loxP PCR product sequencing and alignment

**Development of Novel *Shank*$^{fl/fl}$ Mouse Line**

**A)** Primer design for validating the generation of *Shank3*$^{fl/fl}$ mice. Three primer pairs are designed to target three loxP sites in our previously reported *Shank3 e4-22*$^{flox/flox}$ mouse line.

**B)** Genotype of the founder mouse and its litters after CRISPR/Cas9 engineering. Using the 2$^{nd}$ loxP primer pair, the founder mouse showed 210 bp PCR product while the control mouse showed 320 bp PCR band, suggesting a 110 bp deletion around the 2$^{nd}$ loxP site (Left). Four pups from the founder mouse crossed with WT mouse showed three different genotypes, pup 1 and pup 4 are WT (no PCR band), pup 2 is a *Shank3 e4-22*$^{flox/flox}$ homozygous mouse (showing one 320 bp PCR band), and pup 3 is the *Shank3*$^{fl/fl}$ heterozygous mouse (showing one 320 bp PCR band and one 210 bp PCR band) (Right).

**C)** Sequencing and alignment of the PCR product from three primer pairs. Mouse 1 and mouse 2 are *Shank3 e4-22*$^{flox/flox}$ mouse lines (3 loxP sites). Mouse 3 is a *Shank3*$^{fl/fl}$ mouse (2 loxP sites). Sequence of the loxP site is highlighted with orange color. 110bp deletion, including the loxP sequence, is highlighted in red.

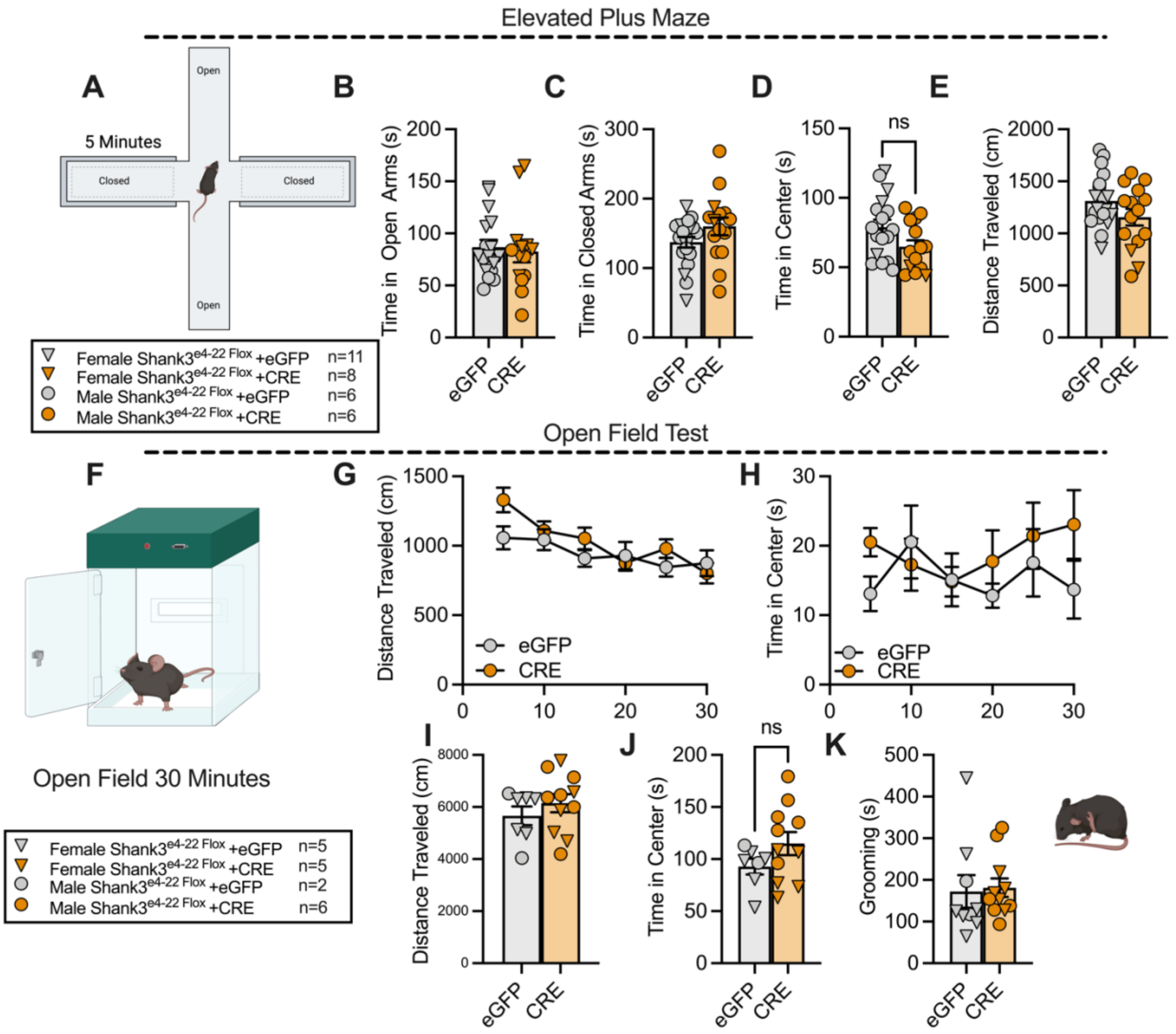


**Figure S2**: **There are no differences in anxiety-like behavior between** ***Shank3*** $^{fl/fl\ +eGFP}$ **and** ***Shank3*** $^{fl/fl\ +CRE}$ **mice**

**A)** Schema of Elevated Plus Maze.

**B-E B)** There were no differences between *Shank3* $^{fl/fl\ +eGFP}$ and *Shank3* $^{fl/fl\ +CRE}$ mice in time in open arms (Unpaired t-test, one-tailed, P=0.3694, *n1= 17*, *n2=14*) **C)** time in closed arms (Unpaired t-test, two-tailed, P=0.1164, *n1= 20, n2=15*) **D)** time in center (Unpaired t-test, two-tailed, P=0.0637, *n1= 17*, *n2=14*) or **E)** distance traveled (Unpaired t-test, two-tailed, P=0.1047, *n1= 20, n2=15*)

**F)** Schema of Extended Open Field Task

**G)** There were no differences between *Shank3* $^{fl/fl\ +eGFP}$ and *Shank3* $^{fl/fl\ +CRE}$ mice in distance traveled across 30-minute test analyzed in 10-minute bins. (Two-way ANOVA, Time x Virus *P=0.022;* $F_{5,80}=2.803$; *n1= 7, n2=11*)

**H)** There were no differences between *Shank3* $^{fl/fl\ +eGFP}$ and *Shank3* $^{fl/fl\ +CRE}$ mice in time in center across 30-minute test analyzed in 10 minute bins. (Two-way ANOVA, Time x Virus *P=0.6367;* $F_{5,80}=0.6367$; *n1= 7, n2=11*)

**I-K) I)** There were no differences between *Shank3* $^{fl/fl\ +eGFP}$ and *Shank3* $^{fl/fl\ +CRE}$ mice in distance traveled (Unpaired t-test, two-tailed, P=0.3631, $n1= 7$, $n2=11$) **J)** time in center (Unpaired t-test, two-tailed, P=0.1645, $n1= 7$, $n2=11$) or **K)** grooming when analyzed in total (Unpaired t-test, two-tailed, P=0.2012, $n1= 8$, $n2=11$).

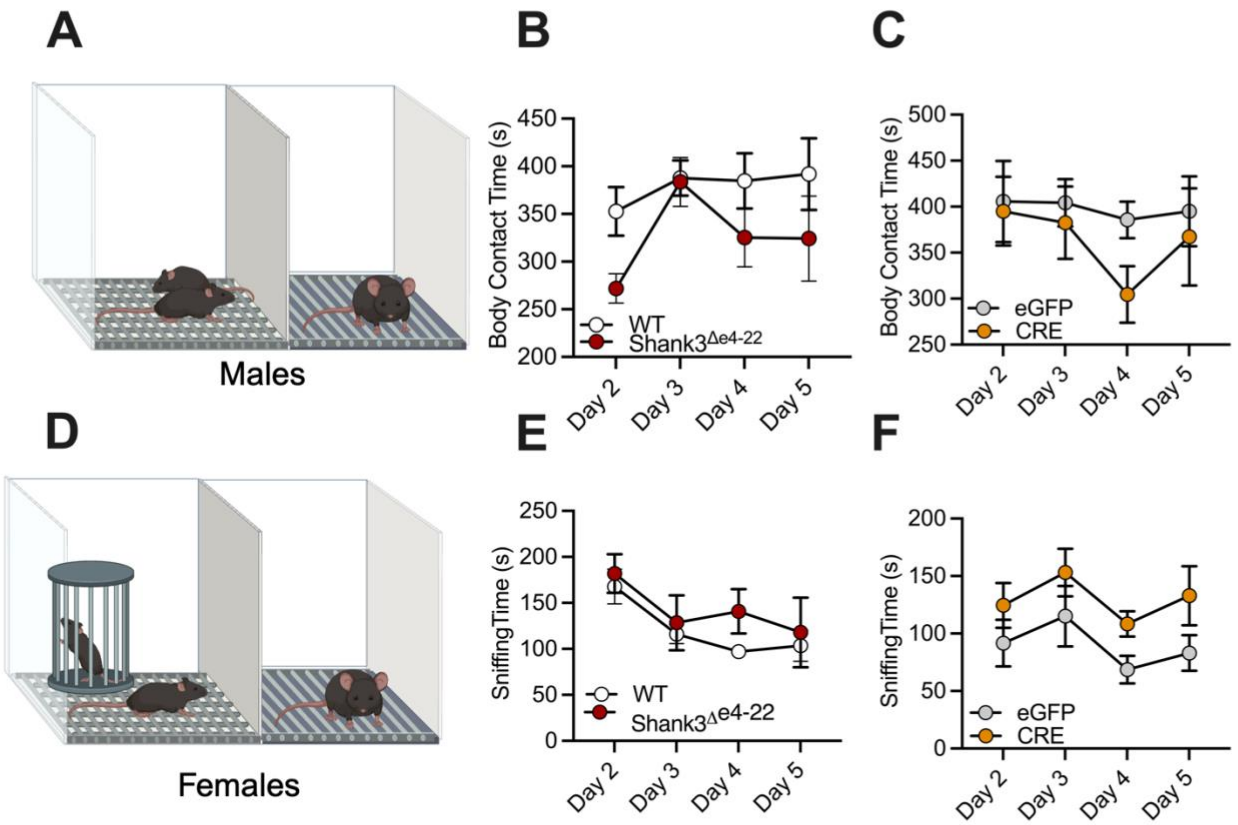


**Figure S3**: **Test groups do not differ in social behaviors during conditioning phase of social conditioned place preference task**

**A)** Schema of conditioning for males.

**B)** No differences were found between *Shank3*$^{\Delta e4\text{-}22}$ and WT mice (Two-way ANOVA, Day x Genotype $P=0.6037$; $F_{3,24}=0.6285$; $n1= 4$, $n2=6$) or **C)** *Shank3*$^{fl/fl\ +eGFP}$ and *Shank3*$^{fl/fl\ +CRE}$ mice in body contact time across the 4 conditioning days. (Two-way ANOVA, Day x Virus $P=0.2804$; $F_{9,54}=1.259$; $n1= 6$, $n2=6$)

**D-F) D)** Schema of conditioning females. No differences were found between **E)** *Shank3*$^{\Delta e4\text{-}22}$ and WT mice (Two-way ANOVA, Day x Genotype $P=0.8537$; $F_{3,31}=0.2598$; $n1= 9$, $n2=6$) or **F)** *Shank3*$^{fl/fl\ +eGFP}$ and *Shank3*$^{fl/fl\ +CRE}$ (Two-way ANOVA, Day x Virus $P=0.9406$; $F_{3,30}=0.1314$; $n1= 7$, $n2=8$) mice in sniffing time during the 4 conditioning days.
